# Cellular consequences, citrullination substrates, and antigenicity resulting from wild-type and targeted PAD4 on cell surfaces

**DOI:** 10.64898/2026.01.05.696859

**Authors:** Sophie Kong, Trenton M. Peters-Clarke, Corleone S. Delaveris, Paul Phojanakong, Veronica Steri, James A. Wells

## Abstract

Protein arginine deiminase-4 (PAD4) catalyzes hydrolysis of arginine to citrulline in proteins that promotes widespread changes in cellular phenotypes through transcriptional regulation that can induce innate immunity and promote cancer. Overexpression and hyperactivity of PAD4 leads to a form of cell death called NETosis that releases PAD4 to the extracellular space. In excess, release of PAD4 is believed to be a major cause of various autoimmune diseases through the generation of anti-citrulline protein antibodies (ACPAs). Little is known about the specific protein substrates that become citrullinated and lead to autoimmunity, but there is growing evidence that PAD4 can be localized to the cell surface in response to inflammation. Here, we further characterize the cellular consequences for exogenous treatment with PAD4 showing that it induces morphological changes that increase cell migration, a hallmark of cancer. We then devised a more simplified and robust proteomics approach to identify PAD4 substrates. We identified some 1000 endogenously citrullinated peptides from 500 proteins, and 3000 citrullinated peptides from 1300 proteins upon exogenous addition of PAD4 both inside and outside of cells. This extracellular set can be further augmented by targeting PAD4 to a cancer target, HER2, using a binding protein conjugate. Finally, we studied how citrullinated cells can induce a robust humoral response in a syngeneic vaccine model to produce ACPAs. We believe these studies further our understanding of cell phenotypic consequences of extracellular PAD4 and new PAD4 substrates both inside and outside of cells that are potential neoepitopes for generation of ACPAs.

## Introduction

Protein arginine deiminase 4 (PAD4) is a calcium-dependent enzyme that catalyzes the conversion of peptidyl arginine to peptidyl citrulline.^1–3^ This post translation modification changes the positively charged guanidium group on Arg to a neutrally charged urea group. Though citrullination results in only a 0.984 Da mass difference, the loss of a positive charge can cause significant structural and compositional changes that induce immunogenicity of substrate proteins.^4–6^ The generation of citrullinated peptides and neo-epitopes is a hallmark in inflammation and a variety of autoimmune diseases. About two-thirds of the patient population in rheumatoid arthritis (RA) develop anti-citrullinated protein antibodies (ACPAs), and these patients tend to exhibit more severe disease phenotypes, suggesting the inflammatory nature of citrullinated neo-epitopes.^7^

The human proteome contains five PAD isoforms, PAD1-PAD4 and PAD6. Of all five, PAD4 is the only isoform that contains a nuclear localization signal and PAD4 is expressed exclusively by immune cells, specifically by monocytes and granulocytes.^8–10^ The intracellular function of PAD4 is to locally citrullinate histones allowing for changes in transcriptional regulation. Rampant induced citrullination leads to a process of inflammatory neutrophil cell death called NETosis.^11–13^ In NETosis, neutrophils burst open and release their intracellular and nuclear contents into the extracellular space, including unravelled networks of DNA and associated proteins forming widespread “nets”. During this process, active PAD4 is known to be carried along the DNA “nets” to enter the extracellular space and synovial fluid, resulting in citrullination of extracellular targets.^14–16^ These substrates are recognized by the immune system, which develops anti-citrullinated protein antibodies (ACPAs) to the neo-antigens and promotes chronic autoinflammation.^12,16^

Recent work from the Durrant group have shown that citrullinated peptides from RA studies can be immunogenic and leveraged to improve immune responses to cancer.^17–20^ Citrullinated peptides have been shown to bind certain human leukocyte antigen (HLA) alleles with higher affinity than their non-citrullinated peptide counterparts, and recognition of these presented peptides by CD4+ T cells elicits an anti-tumor response.^16,18^ When injecting a select combination of citrullinated peptides into tumor-bearing mice, these vaccines have caused tumor regression and production of pro-inflammatory cytokines. These peptide vaccines are composed of citrullinated peptides from vimentin and enolase that are known targets of ACPAs in RA.^21,22^ We hypothesize that there may be many more PAD4 substrates at the cell surface that can be identified and potentially leveraged for similar anti-tumor purposes.

Though canonically an intracellular protein, recent reports have found active PAD4 localized to the cell surface and that its expression levels are correlated with levels of inflammation^23–25^. Here, we study the functional consequences and identify citrullinated products of wild-type PAD4 and a HER2 targeted variant of PAD4 on cells. We develop an improved MS work-flow to identify natural substrates. We show these are broadly pro-inflammatory through *in vivo* vaccination of tumor cells that induces ACPAs in a syngeneic mouse model. Broadly, these studies identify specific extracellular PAD4 citrullinated neo-epitopes that enhance autoreactive antibodies.

## Results

### Visualization of PAD4 on the cell surface

Though PAD4 intracellular activity has been well characterized, we sought to better understand the isolated effects of PAD4 when added exogenously to cells, to mimic the effect of its release from cells undergoing NETosis. To a culture of murine EMT6 breast cancer cells, we added 1 µM of soluble, recombinant PAD4 and imaged every 15 min for 4 hours, then hourly for anther 20 hours. Media was supplemented with 2 mM Ca^2+^ and reducing agent, TCEP, to promote PAD4 activity and stability. Compared to vehicle or a catalytically inactive PAD4 variant (D350A), cells treated with active PAD4 showed a rapid initial decrease in cellular eccentricity within 15 min before returning to normal morphology within 10 hrs (**Fig. 1A**). Eccentricity is a measure of how much the cell deviates from being circular, suggesting that PAD4 treated cells morphologically appear more circular and less adherent than control cells. These changes were also reflected in changes in cell surface area (**Fig. 1B**) where active PAD4 treated cells showed a rapid decrease and then return to initial surface area after 10 hr. The cell counts for vehicle and inactive PAD4 treated cells increased faster than for active PAD4 treated cells, reaching a maximum at 20 hr whereas active PAD4 treated cell numbers gradually increased throughout (**Fig. 1C**). There was little difference between these samples in terms of cell viability with time (**Fig. 1D**).

**Fig. 1.**
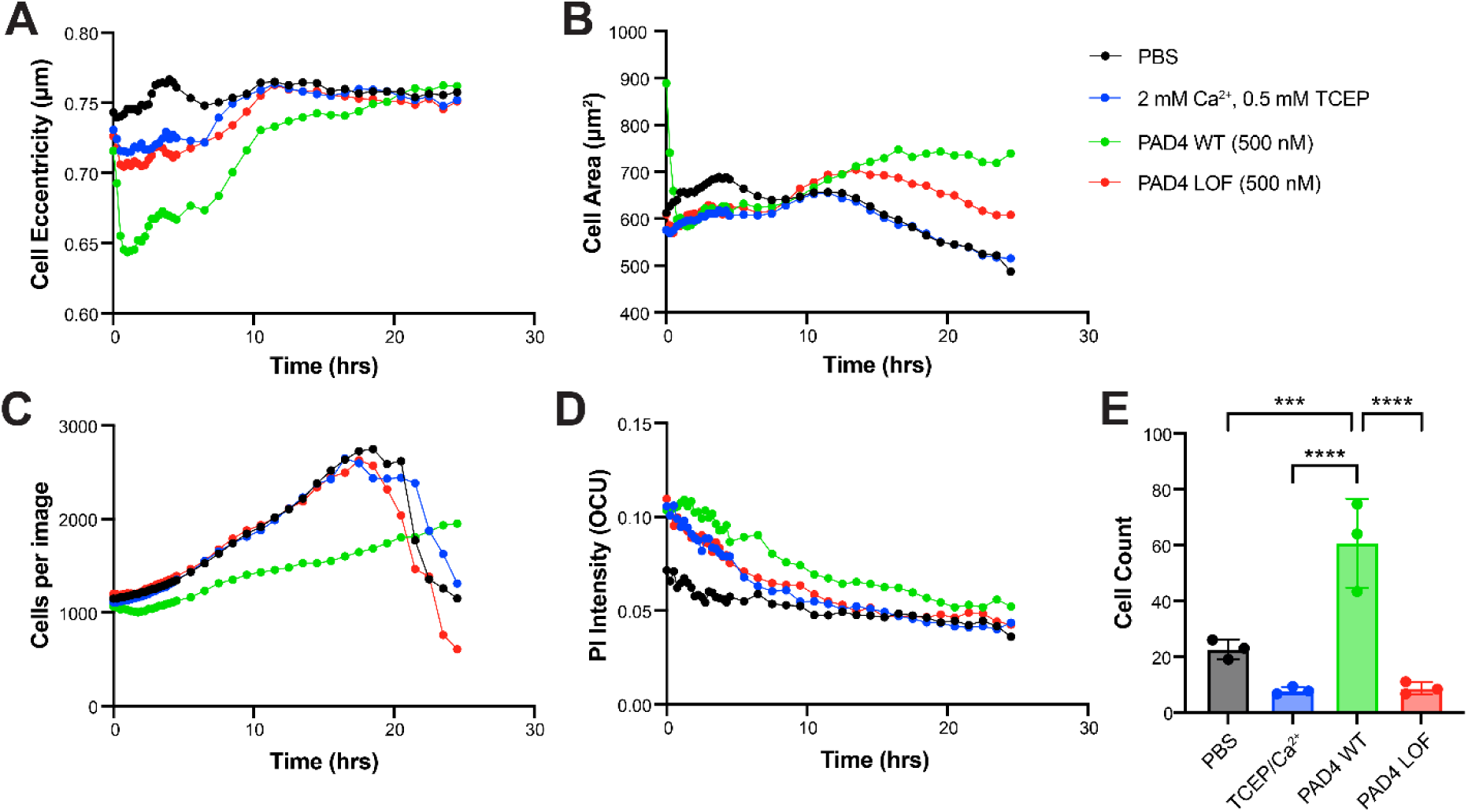
Treatment of EMT6 cells with active PAD4 versus a loss-of function variant differentially changes cell morphology, growth rates, viability, and migration. Cells were treated for 24 hours and different parameters measured by incucyte. (A) Cellular eccentricity defined as a less spherical cell shape, (B) cell area, (C) cell count, and (D) cell viability measured by propidium iodine stain. (E) Transwell migration of Patu8988t cells across a porous membrane. PAD4 treated cells have an increase propensity for migration as compared to untreated, vehicle, or PAD4 LOF treated cells.

### Migration of PAD4 treated cells

PAD4 has been implicated in promoting cancer metastasis.^6,26,27^ Our incucyte data showed that the morphology of PAD4-treated cancer cells remarkably mimics the morphology of cells treated with versene, a reagent commonly used in cell culture to lift adherent cells from cell culture flasks. This suggested that PAD4 treatment of cells leads to disruption of cell adherence that can serve as a precursor to migration. To test if PAD4 treatment would enhance cell migration, we used a transwell migration assay (**Fig. S1**). Patu8988t cells, which show low levels of cellular migration when untreated, were plated in serum-free media, and we monitored migration into complete media, plus or minus PAD4 or variants. After 24 hours, cells treated with PAD4 showed significantly increased migration to the serum-positive media as compared to the untreated or PAD4 D350A inactive controls (**Fig. 1E**).

### Engineering of a PAD4 targeted to HER2 expressing cells

Tethering post-translationally modifying enzymes, such as kinases or subtiligase, to the surface of cells can enhance their surface modifying activities by over ten-fold.^28–32^ Thus, we wanted to test if this were the case with PAD4 if we could target it specifically to HER2 over-expressing cells.^33^ The therapeutic HER2 antibody trastuzumab has been extensively used to direct small-molecule cytotoxins, peptides, and enzyme-based payloads selectively to HER2-expressing cells.^34–39^ Several attempts were made at expressing a trastuzumab IgG or Fab conjugate with PAD4, but genetic fusions suffered from low expression yields and chemical conjugates led to protein instability and aggregation. This was not surprising as PAD4 contains numerous surface-exposed cysteines and requires storage with reducing agent to prevent aggregation while trastuzumab has stabilizing disulfides which makes their expression as a fusion very challenging.

ZHER2 is a small three-helical bundle affibody protein, derived from the Z domain of bacterial Protein A that contains no disulfide bonds.^40^ The affibody is engineered to bind tightly to HER2 with a Kd of 22 pM.^40^ ZHER2 has even been used clinically as a PET imaging diagnostic with reduced immunogenicity considerations.^41,42^ Thus, we engineered a ZHER2-PAD4 fusion containing an N-terminal His-Avi-TEV sites for purification, immobilization, and cleavage, followed by ZHER2 and a 10 amino acid glycine/serine linker to PAD4 (**Fig. 2A**). The ZHER2-PAD4 fusion protein was successfully expressed in *E. coli* and retained binding to the HER2 ectodomain as measured by biolayer interferometry (BLI) (**Fig. 2B)**. We next compared the ability of the ZHER2-PAD4 versus wild-type PAD4 to bind to HER2 over-expressing cells via flow cytometry (**Fig. 2C**).

**Fig. 2.**
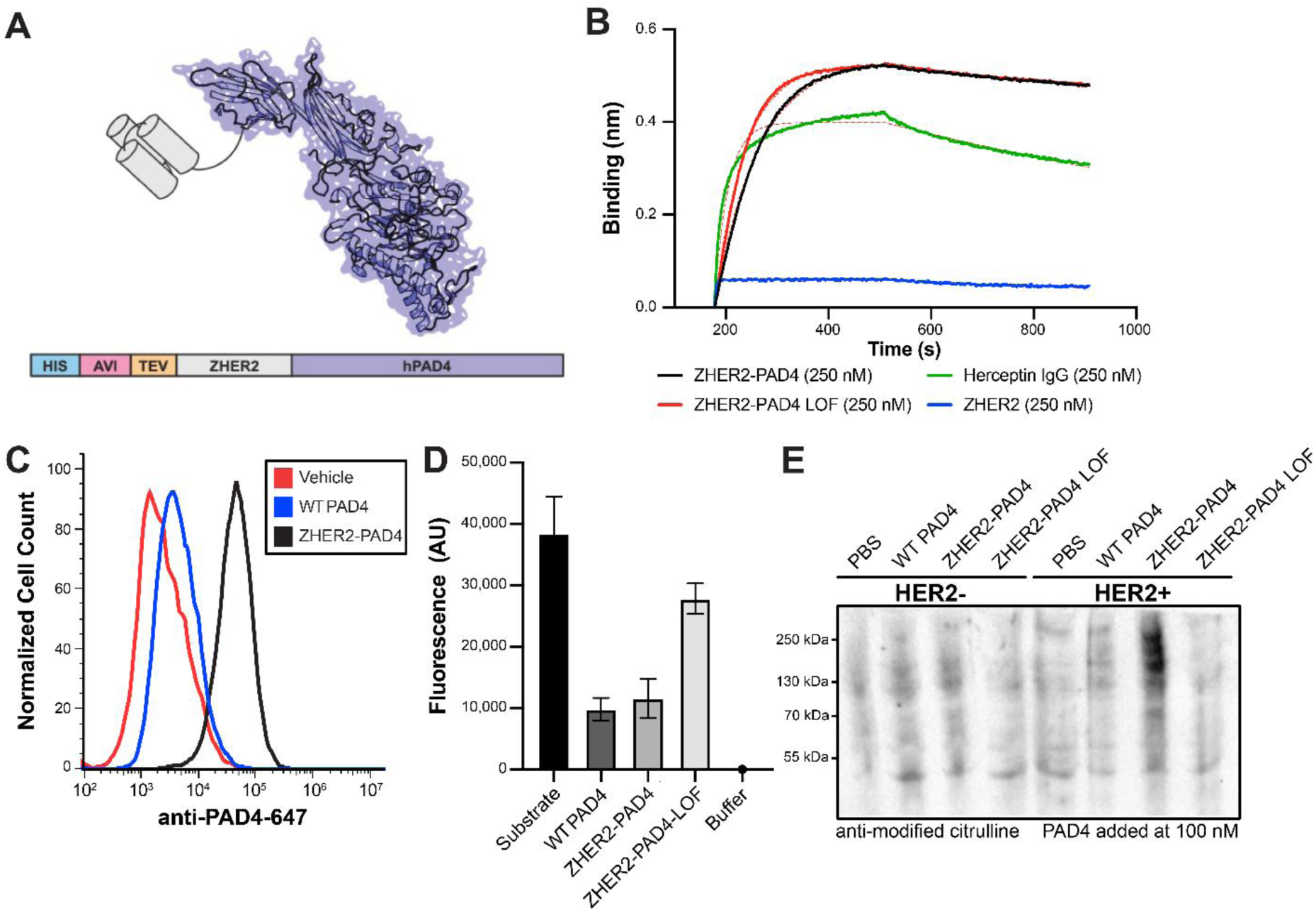
Design and characterization of a ZHER2-PAD-4 fusion for targeting HER2 expressing cells. (**A**) Schematic representation of ZHER2-PAD4 fusion with N-terminal His and Avi tags for purification or immobilization followed by a TEV cleavage site for convenient removal of the tags. (**B**) Binding of ZHER2-PAD4, a loss-of-function (LOF) variant, the HER2 Fab derived from trastuzumab, or the ZHER2 nanobody to HER2 ectodomain measured by bi-layer interferometry (BLI). All bind with similar affinities and in proportion to molecular weight as expected. (**C**) Binding of 100 nM ZHER2-PAD4 or wild type (WT) PAD4 to EMT6 HER2+ cells measured via flow cytometry. (**D**) Activity of PAD4 variants as measured by self-quenched fluorescent trypsin substrate. Note that both WT and ZHER2-PAD4 citrullinate the substrate thus blocking trypsin hydrolysis and reducing fluorescent signal. (**E**) A western blot showing generation citrullinated proteins generated in EMT6 wild type (left) or transfected with HER2 (right) upon treatment with PBS, WT PAD4, ZHER2-PAD4, its LOF variant. Note the increased levels of PAD4 citrullination activity and the dramatic increase upon tethering by ZHER2-PAD4 to HER2 containing cells.

Cells were incubated with each PAD4 variant and then stained with an anti-PAD4-Alexa647 fluorescent probe. Interestingly, the wild-type PAD4 showed some binding relative to untreated control but the ZHER2-PAD4 showed dramatically increased binding. The activity of PAD4 in the ZHER2-PAD4 fusion was comparable to wild-type PAD4, as measured by detecting citrullination of an arginine-mimetic pro-fluorescent substrate (**Fig. 2D**). In this assay, a self-quenched substrate is incubated with PAD4. If the substrate is citrullinated it cannot be cleaved by trypsin and remains self-quenched. Thus, the level of fluorescence after treatment with trypsin is inversely proportional to PAD4 activity. This HER2 targeted form of PAD4 is also more active at the surface of HER2 expressing cells compared to inactive controls and untargeted, WT PAD4 (**Fig. 2E**).

### Identifying sites of citrullination using isocyanate triggered ETD proteomics

The small difference in mass between Arg and citrulline (<1 Da) presents a challenge for confident identification of sites of citrullination. Further, this mass shift is nearly isobaric with the peptide’s isotopic envelope, necessitating high resolution mass analysis and obfuscating downstream spectral interpretation.^43^ Two proteomic methods are conventionally employed to address these challenges. The first involves conjugation of citrullinated peptides with citrulline-specific enrichment handles followed by liquid chromatography (LC) fractionation of tryptic digests.^44,45^ However, this method is time consuming and requires harsh conditions for peptide labelling. A second strategy involves a combination of higher energy collisional dissociation (HCD, aka beam-type collision induced dissociation (bCID))^43^ and electron-transfer dissociation (ETD) of the entire tryptic digest to identify sites of citrullination. ETD enables indiscriminate peptide backbone fragmentation without loss of labile post-translational modifications (PTMs)^46–51^, but is time consuming and subjecting every precursor to ETD reduces the throughput and depth of analysis.^43,52^ We implemented an alternative data acquisition scheme that relies on HCD fragmentation of peptidyl citrulline to induce release of the isocyanate group (**Fig. 3A**).^53,54^ Only upon detection of this 43 Da neutral loss does the mass spectrometer trigger an electron-transfer/higher-energy collision dissociation (EThcD) scan of the corresponding precursor, greatly reducing the overall duty cycle (**Fig. 3B**).^55^ This method is referred to as HCD-product dependent-EThcD (HCD-pd-EThcD). By manually inspecting spectra from the triggered EThcD scans, sequence-informative b, c, y, and z• ions allow confident identification of citrullinated peptides and localize the 0.984 Da shift to the appropriate Arg residue (**Fig. 3D**). We validated this approach on a simple *in vitro* system containing only recombinant PAD4 and histone H3, a well-known substrate. PAD4 and histone H3 were incubated together at 37°C for 90 minutes in the presence of 0.5 mM TCEP and 2 mM Ca^2+^ for optimal PAD4 activity to allow citrullination prior to trypsin digestion for proteomic analysis. The PBS and inactive PAD4 control conditions were subjected to the same workflow. We generated complete sequence of the 136 amino acid histone H3 protein and identified 12 unambiguous sites of Arg citrullination, all of which matched published reports (**Fig. 3E**). Previous research identified four additional citrullination sites on H3 using a partial proteolysis strategy to cover Arg-dense regions.^56^

**Fig 3.**
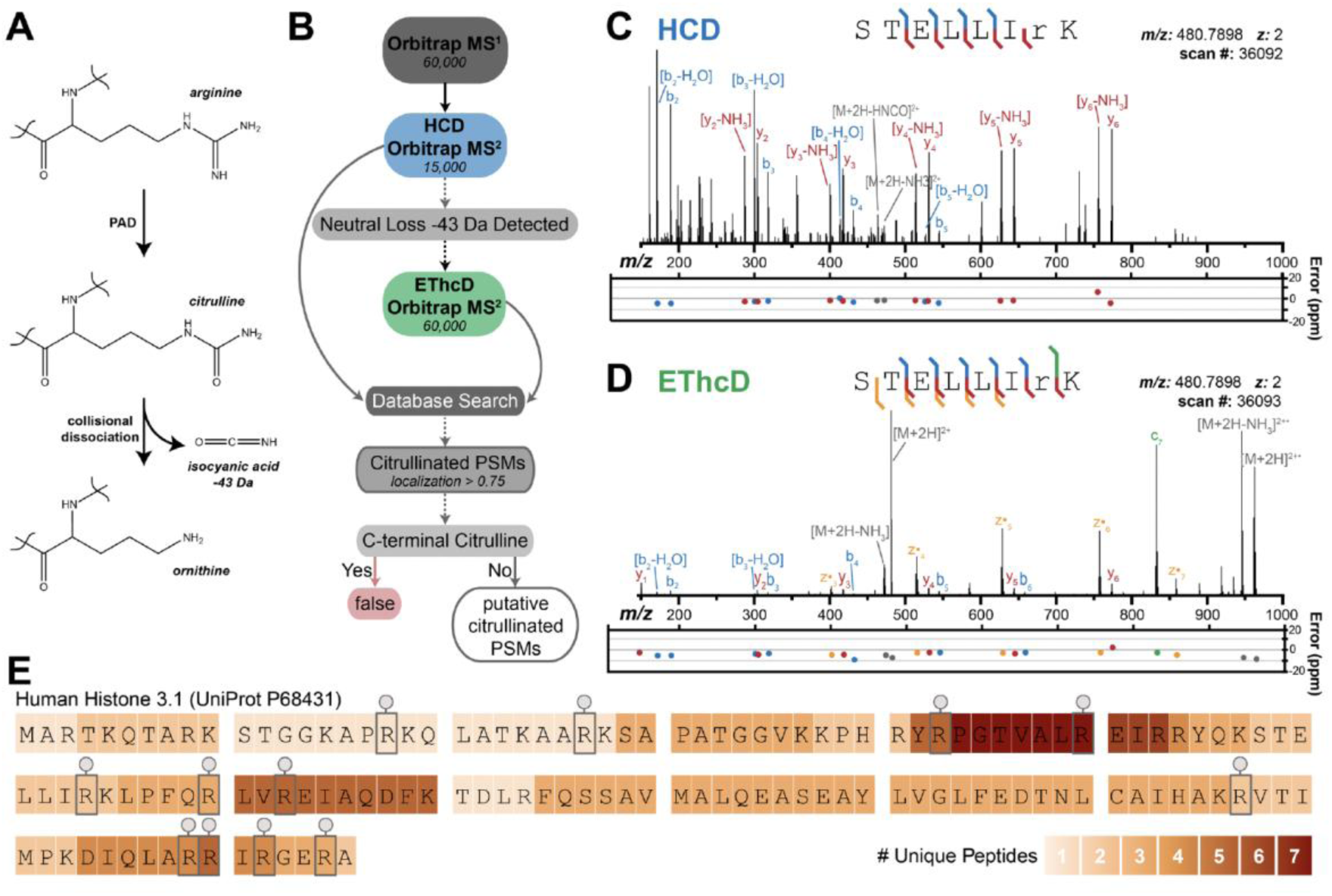
Mass spectrometry workflow and validation using recombinant PAD4 and a well-established PAD4 substrate, histone H3. (**A**) Chemical structures for the generation of citrulline from arginine and the dissociation of citrulline to ornithine upon collisional activation that yields neutral loss of isocyanate. (**B**) Proteomic data acquisition and spectral analysis workflow used to identify and filter citrullinated peptides. Performing high-energy collisional dissociation (HCD) coupled to product-dependent, triggered electron transfer/higher energy collisional dissociation (EThcD) scans (HCD-pd-EThcD) upon detection of isocyanic acid loss from the precursor, provided an optimized method for higher throughput and accuracy of detecting citrullinated products. An example of the two steps (**C**) HCD followed by (**D**) EThcD annotated spectrum for the histone H3 peptide STELLIRK (amino acids 58-65) containing a site of citrullination.^76^ (**E**) Twelve high-confidence sites of arginine citrullination mapped onto the amino acid sequence of histone H3 using HCD-pd-EThcD. Protein coverage is displayed in a heatmap, illustrating 100% sequence coverage.

### Identifying sites of citrullination by exogenous addition of PAD4 to whole cell lysates

Following optimization *in vitro* to identify citrullination events, we moved to a more complex sample of whole cell lysate treated with PAD4. A human HER2+ expressing cell line was generated by transducing EMT6s murine breast cancer cells with a hHER2 expressing lentiviral vector. Cells were lysed and treated with PBS, PAD4, PAD4 D350A or ZHER2-PAD4, then digested and fractionated offline by HPLC to ensure greater separation of peptides. Samples were prepared for downstream proteomic analysis using the HCD-pd-EThcD method. In each sample, about 6,000 proteins were identified from close to 50,000 total peptides. In PAD4 active samples, we identified more than 3,000 unique citrullinated peptides and 1,300 proteins, whereas PBS and inactive PAD4 treated control samples each yielded only about 1,000 citrullinated peptides and 500 proteins (**Fig. 4A-C**). To reduce false positives in the dataset we further filtered out peptides with citrulline localization confidence below 75%. The activity of trypsin to cleave after citrulline is reduced by >100 fold compared to Arg; such events would leave C-terminal citrulline. To remove these rare artifacts, we filtered out all identified peptides containing C-terminal citrullines.^55,57,58^ This filter reduced the number of citrullinated peptides detected in negative control samples (∼30%) while minimally affecting our active PAD4 datasets. Analysis of unique citrullinated proteins shows that most citrullinated proteins observed in PBS or PAD4-LOF treated conditions are also found in active PAD4 treated conditions as expected since these samples would also include endogenous citrullination (**Fig. 4D**). Interestingly, half of citrullinated proteins identified upon treatment with active PAD4 were modified more than once, consistent with the heightened activity for exogenous PAD4 treatments (**Fig. 4D, E**).

**Fig 4.**
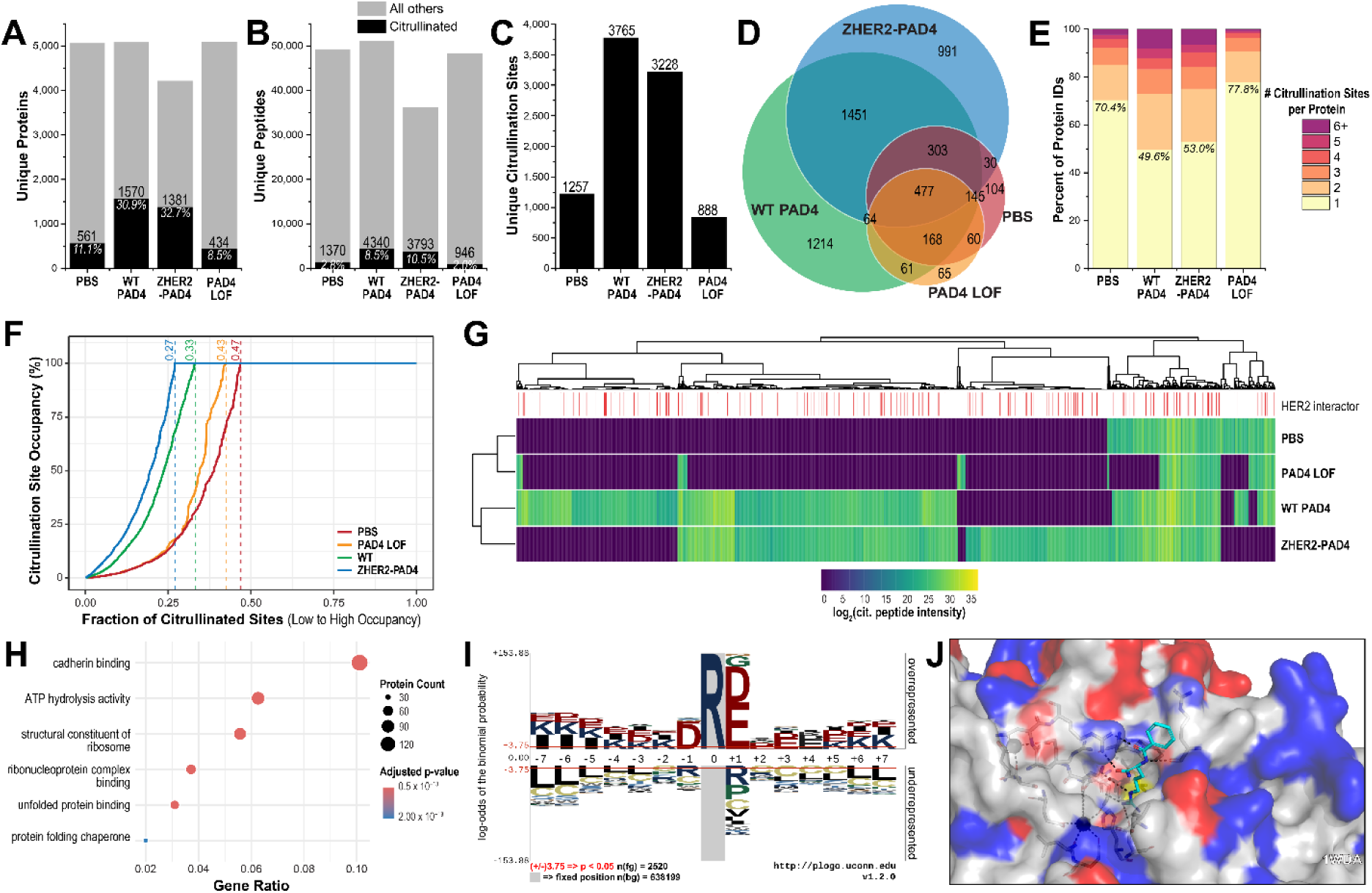
Deep proteomic profiling of cell lysate samples treated with PAD4 to identify unique, citrullinated substrates. Number of citrullinated (**A**) proteins, (**B**) peptides, and (**C**) sites identified from the four treatment conditions along with all non-citrullinated sites. (**E**) Number of citrullination sites identified per protein. Note that treatment with active PAD4 produced 3 to 4 times more citrullinated peptides, proteins, and citrullination per protein than conditions without exogenously added PAD4 or PAD4 LOF. (**F**) Citrullination site occupancy as a cumulative distribution plot for each treatment condition. For each site of high-confidence citrullination, the intensity of citrullinated site was compared to the intensity of the unmodified site, including multiple isoforms and charge states, to reach saturation. Each condition was rank ordered by increasing citrullination site occupancy. Fractions along the top of the plot indicate the fraction of all citrullination sites within that condition which have less than 100% site occupancy. (**G**) Global heat map analysis of citrullinated peptides from each experimental group with HER2 interacting proteins highlighted in red. Protein–protein interaction data for HER2 were obtained from BioGRID.^60^ (**H**) Gene ontology and molecular function analyses of proteins that were elevated in the ZHER2-PAD4 treatment of EMT6 lysates. (**I**) Consensus sequence Logo of citrullinated peptides. PAD4 shows strong preference for negatively charged residues surrounding site of citrullination and other charged residues on the flanks. (**J**) Citrullinated peptide analogue bound at the active site of PAD4 enzyme (PDB: 1WDA). Several basic residues (blue) on PAD4 shown to interact with flanking negatively charged residues on citrullinated peptide.

Examination of the site occupancy of citrulline at endogenous Arg residues across the proteome again reveals the broad activity of PAD4 (**Fig. 4F**). Each localized site of citrulline was assessed an occupancy rate from 0 to 100% based on the intensities of modified and unmodified peptide isoforms capturing that residue. By monitoring the extent of citrullination at each site and rank ordering the citrullination sites from lowest to highest occupancy, the activity of exogenous PAD4 is seen. Of the 1257 and 888 unique citrullination sites detected in PBS and PAD4-LOF treatments of EMT6 lysate, respectively, just over half of these sites are fully occupied by citrulline. Conversely, active PAD4 treated lysate yielded 3,765 unique citrullination sites with essentially complete occupancy at 67% of all sites. The ZHER2-PAD4 treated lysate gave a slightly lower 3,228 unique citrullination sites, but a higher proportion (73%) of those sites were completely occupied. Of note, these calculations estimate site occupancy based on peptide label-free quantification (LFQ) intensities; they do not consider differences in electrospray ionization or lack of coverage of the unmodified peptide counterparts. We believe these metrics provide valuable insights into the breadth of PAD4 activity.

Of the top 150 proteins identified in the exogenous active PAD4 datasets, dozens of membrane or membrane-associated proteins stood out to us, including HER2 and transferrin receptor (**Table S1**). We generated a global heat map of all proteins identified to be citrullinated, separated by experimental condition (**Fig. 4G**). Though there is a large overlap of protein substrates between WT PAD4 and ZHER2-PAD4 treated samples, each condition also contains a unique set of citrullinated proteins. When looking at proteins only citrullinated by ZHER2-PAD4, we do not observe any trend in there being more known HER2 interactors (**Fig. S2**). This likely reflects how abundant HER2 is on the surface of cells and how PAD4 is a so promiscuous that citrullination can occur rampantly on any membrane protein whether cells are treated with WT PAD4 or ZHER2-PAD4. Indeed, most of the upregulated citrulline sites upon ZHER2-PAD4 treatment relative to PBS and WT PAD4 are known HER2 interaction proteins. Proteins that were upregulated for citrullination in ZHER2-PAD4 treatment were used for gene ontology (GO) term enrichment analysis. Molecular function analysis revealed proteins associated with cadherin binding, ATP hydrolysis activity, structural constituents of the ribosome, unfolded protein binding, and protein folding chaperones (**Fig 4H**). Cellular component analysis revealed proteins associated with the cell-substrate junction and focal adhesion among the ZHER2-PAD4 citrullinated targets, signifying some enrichment for the cell surface substrates, even within cell lysate (**Fig S7-8**). We also identify ZHER2-PAD4 substrates canonically annotated as intracellular proteins. The abundances of these intracellular sites are often not significantly different from WT PAD4 levels, indicating that active PAD4’s promiscuity and broad substrate scope govern modification of these intracellular proteins with EMT6 lysate, not driven by induced proximity through HER2 binding as is the case with true HER neighbors on intact cells.

Previous studies using histone H.3 peptide derivatives have shown PAD4 to be a promiscuous enzyme with broad substrate specificity centered around the Arg.^59^ Our studies corroborate these findings. We do find a mild consensus sequence that slightly favors acidic residues, along with glycine, directly before and after the site of citrullination (**Fig. 4I**). This substrate specificity can be explained by inspection of the substrate binding site of PAD4 (PDB ID: 1WDA). Multiple basic residues reside in the substrate binding pocket of PAD4 that may interact with the acidic residues flanking the citrullinated Arg (**Fig. 4J**).

### Identifying sites of citrullination on the HER2 ectodomain

Through treating lysates with PAD4 and ZHER2-PAD4, we were able to identify unique proteins in each of our data sets. When comparing ZHER2-PAD4 treated lysate to PBS treated or WT PAD4 treated, we find that many proteins including HER2 are vastly enriched in the ZHER2-PAD4 sample (**Fig. 5A**) Annotating previously reported interacting partners of hHER2 reveals that many of the citrullination sites significantly upregulated in the ZHER2-PAD4 treatment are within neighbors or direct interactors of HER2.^60^ We repeated this study using whole SKBR3 cells that express high levels of HER2, treating again with PBS, WT PAD4, ZHER2-PAD4, or PAD4 LOF.^61,62^ These cells were treated *in situ,* then the membrane proteins were isolated using a subcellular fractionation kit. The membrane proteins were then digested and analyzed via the previously described HCD-pd-EThcD workflow.

**Fig. 5.**
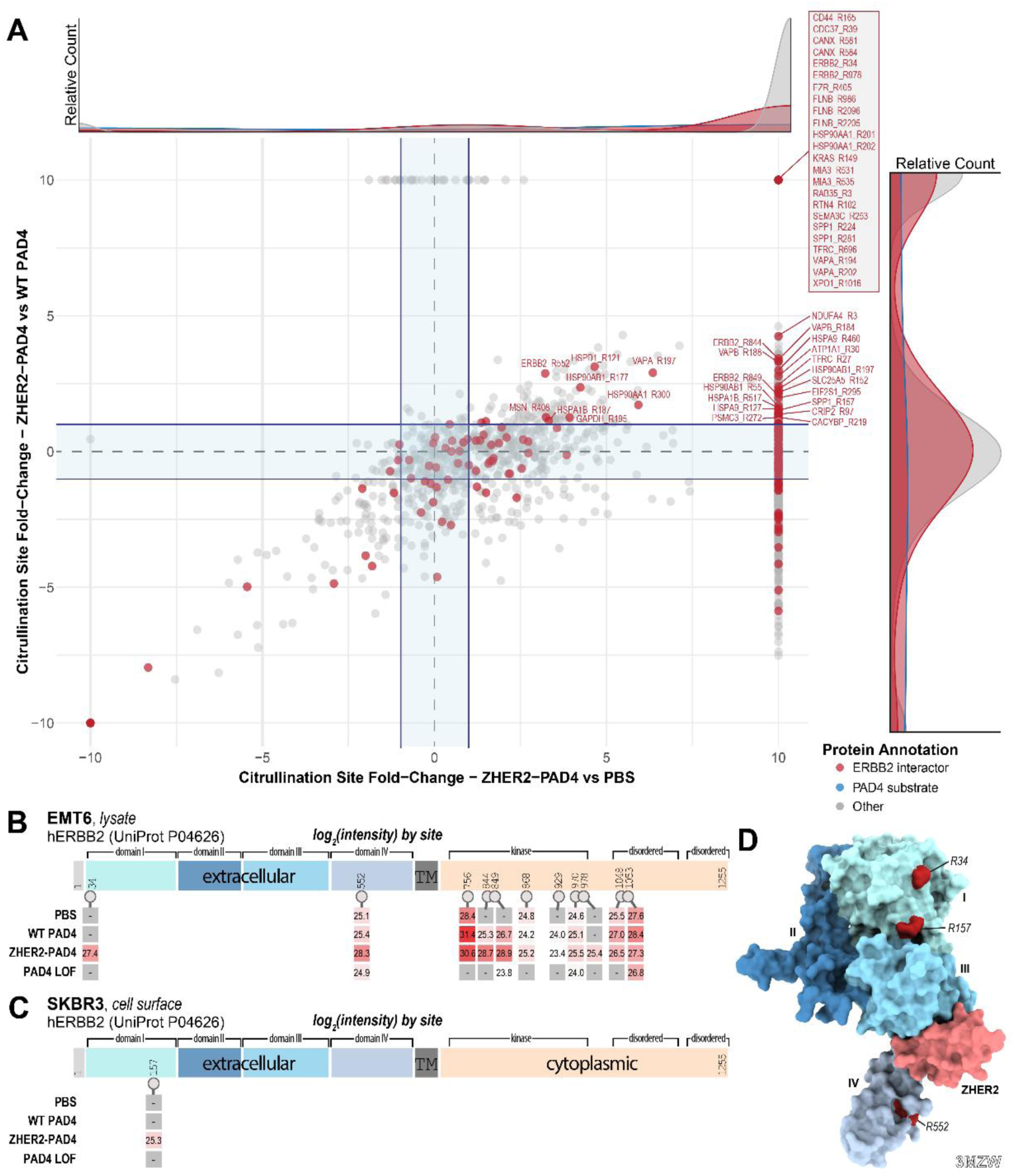
Characterization of citrullinated proteins from PAD4-treated samples and mapping sites of citrullination onto HER2 (hERBB2). **(A)** Enrichment of citrullinated sites identified in ZHER2-PAD4 treated hHER2-ETM6 lysate plotted relative to endogenous citrullinated proteins identified in PBS treated (*x-axis*) or WT PAD4 (*y-axis*) treated lysates. Diagonal shows there is a general correlation for ZHER2-PAD4 activity whether compared to WT PAD4 or to PBS treatments. Sites within the center of the plot reflect mostly non-specific citrullination by PAD4. Many sites of citrullination shown at the far right, including HER2 sites, are found exclusively in the ZHER2-treated samples compared to PBS treatment. The upper right quadrant highlights sites that are heavily enriched in the ZHER2-PAD4 group relative to both background controls. **(B)** Mapping sites of hHER2 citrullination and their abundance from PAD4-treated EMT6 lysate. More sites of citrullination were identified in active PAD4 conditions and with higher abundance. In lysates both intracellular and extracellular sites of citrullination were observed; however, the differential signal for treatment with ZHER2-PAD4 vs PBS was higher for extracellular proteins. **(C)** Mapping sites of hHER2 citrullination and their abundance from PAD4-treated SKBR3 cells. The extracellular R157 site is the only site identified to be citrullinated in the ZHER2-PAD4 condition and no other condition yields hHER2 citrullination. **(D)** Space filling rendering of the extracellular domain of hHER2 (showing the four sub-domains in different colors) bound to the ZHER2 binder in pink (PDB: 3MZW).^40^

In both the EMT6 lysate and SKBR3 cell surface groups, HER2 was found to be citrullinated most by the HER2 targeted ZHER2-PAD4 fusion protein (**Fig. 5B,C**). We mapped sites of citrullination onto the sequence of HER2 and found that while there were many citrullinated peptides identified in the HER2 cytoplasmic domain from both endogenous and exogenous PAD samples, only the ZHER2-PAD4 treated samples produced citrullinated peptides on the HER2 (**Fig. 5B, C**). Lastly, we mapped these extracellular citrullination sites to the structure of ZHER2-bound HER2 ectodomain. ZHER2 binds HER2 between domains III and IV, and all three identified sites of citrullination lay adjacent to the ZHER2 binding domain supporting localized citrullination (**Fig. 5D**).

### Humoral response to citrullinated cell surfaces

Since citrullination is known to be an immunogenic PTM, we investigated if citrullination onto tumor cell surfaces would generate immune responses to cancer cells *in vivo.* We chose EMT6 cells, a syngeneic mouse cell line in the middle of the spectrum between hot and cold tumors.^63^ This cell line has previously shown moderate immune cell infiltration, so we believe that generating immunogenic neo-epitopes could take advantage of tumor infiltrated lymphocytes (TILs) to improve immune response.

As a proof-of-concept experiment, we targeted PAD4 to EMT6 cells *in vitro* to generate cell surface citrullination and then heat-shocked the cells to prevent further cell biology. Also, mild heat shock of cells has been shown to act as an adjuvant for T cell infiltration of tumors, so we decided to use this in combination with cell surface citrullination to create a whole cell vaccine to probe the humoral immune response towards citrullination.^64,65^ BALB/c mice were vaccinated with citrullinated and heat-shocked whole cells on a weekly basis for four weeks on alternating flanks. Following four weeks of vaccination, WT EMT6 cells or citrullinated EMT6 cells treated with PAD4 were implanted orthotopically to study tumor take and growth rate (**Fig. 6A**). Five mice were used for each condition, and at the end of four weeks, mice were sacrificed and serum was isolated from the blood.

**Fig 6.**
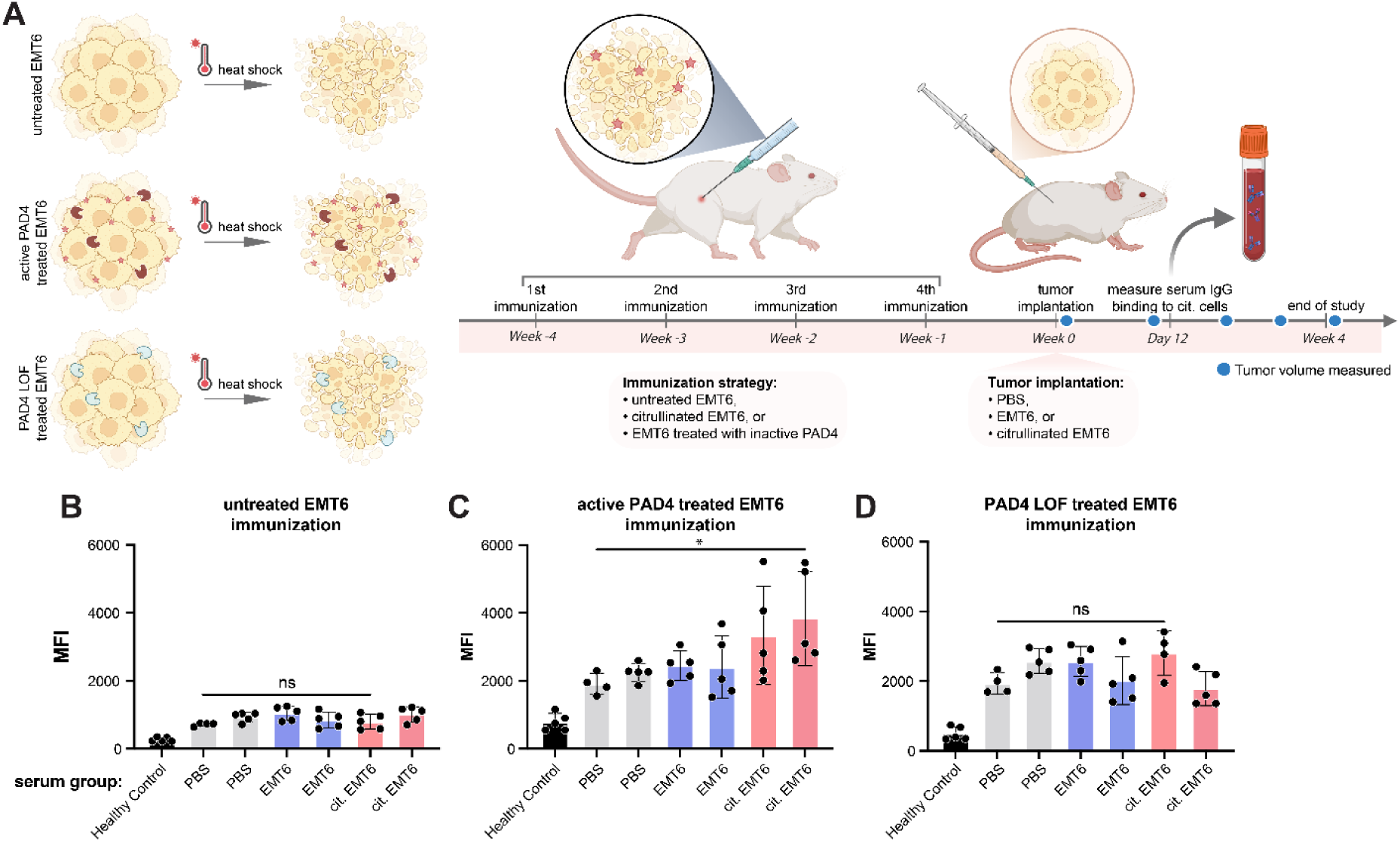
Serum harvested from immunized mice show increased anti-citrullination antibody response to EMT6 cells treated with WT PAD4 versus LOF or PBS control. (**A**) Balb/C mice were immunized 4x on a weekly basis with mouse EMT6 cells that were treated either with PBS, a LOF PAD4, or WT PAD4 to induce extracellular citrullination. EMT6 cells were implanted orthotopically and tracked for tumor growth. Flow cytometry analysis of serum IgG binding to citrullinated EMT6 cells using serum from mouse immunized with (**B**) untreated EMT6 cells, (**C**) active PAD4 treated EM6 cells, or (**D**) PAD4 LOF treated EMT6 cells. Serum was collected for analysis 12 days post implantation with PBS, untreated EMT6 cells, citrullinated EMT6 cells, or from healthy controls. Sample groups were split into 3 pools based on immunization strategy: untreated EMT6, PAD4 treated EMT6, and PAD4 LOF treated EMT6 cells. Each cell population was incubated with serum from immunized mice (via PBS, EMT6, cit. EMT6) or healthy control, separated as groups of 5 mice into 7 different treatment groups (*x-axis*). Created in https://BioRender.com.

To test for an antibody serum response, cells were citrullinated *in vitro* and incubated with serum from the cell vaccinated mice. Binding of mouse IgGs to citrullinated cells was detected via flow cytometry using a fluorescent anti-mouse IgG secondary (**Fig. 6B**). In the group of mice vaccinated with citrullinated cells, serum from several mice showed serum binding to citrullinated cells, while mice vaccinated with wild-type (WT) EMT6 or EMT6 cells treated with inactive PAD4 D350A did not exhibit serum binding to citrullinated cells. This indicates that mice vaccinated with citrullinated cells generated a humoral response specifically to the citrullinated neo-epitopes (**Fig. 6B**). Though a humoral response was generated, there was no observed difference in tumor take rate (**Fig. S3**). We hypothesized that mice that have been vaccinated with citrullinated cells may experience a slower tumor growth rate since their immune systems have been trained to attack citrullinated cells. However the lack of difference in tumor growth rate can be attributed to many factors, one being that syngeneic EMT6 cells grow extremely aggressively when implanted orthotopically. It is possible that the fast-growing nature of the cells completely overshadowed the immune response developed against citrullinated cells.

## Discussion

PAD4 is one of the most highly correlated proteins in RA and has also been associated with cancer and metastasis.^6,8,13,23,26,27^ While its normal function is as a chromatin remodeler via histone citrullination, it also serves a key role in the unspooling of DNA during the process of macrophage cell death.^9,26^ During this process, PAD4 is released to the extracellular space where its hydrolytic activity can citrullinate extracellular proteins and cells in ways that are poorly understood.^9,12,26,61^

Here we show that addition of recombinant PAD4 had dramatic effects on cell morphology and migration that was dependent on its citrullination activity. In fact, these same effects can be phenocopied by addition of EDTA that removes di-cations that reduces cell adhesion. Conversion of Arg to citrulline would reduce the cationic nature of cell surface proteins.^16^ Indeed, we found that active PAD4 treatment increases cell migration and may help to explain how PAD4 can act as a tumor and metastasis promoter.

We next wished to study the mechanism and protein substrate specificity of exogenous PAD4 for both total and cell surface proteins. We were surprised to see that WT PAD4 has natural affinity for cell surfaces as seen by flow cytometry. We and others have shown that when targeted to cell surfaces, enzymes that induce post-translational activities such as glycosidases, kinases, and peptide ligases, can dramatically increase the surface-directed activities by 2-3 logs.^28–32^ We found it is possible to augment this cell surface binding property of PAD4 by fusing it to ZHER2, a small affibody that targets HER2. This further enhanced the binding of PAD4 to cell surfaces for cells expressing HER2 as well as the citrullination of cells as monitored by flow and western blot.

Several proteomic studies have looked at the specificity of PAD4 for predominantly intracellular events by installing covalent handles for citrulline, or by using an exhaustive mix of collisional dissociative events using tandem HCD/ETD on all tryptic peptides.^44,66,67^ Here, we developed and validated a more efficient HCD-pd-EThcD proteomic method that triggers analysis of peptide precursors by EThcD for only those that release ∼43 Da isocyanic acid upon HCD, a characteristic neutral loss of citrulline.^53^ We identified many new endogenous citrullinated substrates and sites that are unique to exogenously added active PAD4. In addition to HER2, we observed citrullination of various cell surface proteins, including CD44 (**Fig. 5A**). CD44 is a cell surface receptor known for its role in cell adhesion and migration.^68–70^ Citrullination of this protein may contribute to the increase migration rate observed with citrullinated cells compared to non-citrullinated cells.^68,69^ Interestingly, citrullinated CD44 peptides were found more enriched in ZHER2-PAD4 cell treated conditions than WT PAD4 treated conditions. Looking at the consensus sequence of all citrullinated PAD4 substrates, PAD4 shows moderate specificity for charged residues, especially acidic residues, flanking the central Arg target. This aligns with charged residues surrounding the catalytic cysteine in PAD4.

Given that the extracellular activity of PAD4 is thought to generate ACPAs, we were very interested in the surface proteins that were citrullinated.^7,13,23^ Indeed, we identified more than 2600 surface proteins that were citrullinated by exogenously added PAD4 (WT or ZHER2-PAD4) not seen in PBS or inactive PAD4 control conditions. We also showed a significant overlap (1,451 sites) with ZHER2-PAD4 as well as 991 substrate sites that were unique to ZHER2-PAD4. Gene ontology and molecular function enrichment analysis reveals these substrates unique to ZHER2-PAD4 treatment include proteins involved in cadherin binding, ribosome biogenesis, unfolded protein binding, and ATP hydrolysis. Moreover, analyzing the lysate of murine EMT6 cells and the cell surface proteins of SKBR3 cells, we find several residues of the hHER2 ECD with citrullination highly upregulated by ZHER2-PAD4. Further, while extracellular and cytoplasmic hHER2 sites are citrullinated by WT PAD4 and ZHER2-PAD4 in EMT6 lysate, treatment of intact SKBR3 cells yields citrullination of only extracellular HER2 arginines and only for the ZHER2-PAD4 condition. Notably, the HER2 binding domain of ZHER2 and the tether length between ZHER2 and active PAD4 enzyme dictate the residues of hHER2 or proximal neighbors on the cell surface that may be citrullinated. Therefore, backed by our results of targeted hHER2 citrullination, we posit that this strategy of targeting a promiscuous, PTM installer or remover enzyme to a protein of interest may be a viable, enzymatic route of proximity labeling proteomics.

Given the importance of PAD4 in RA, we wondered about the immunogenicity of citrullination on cells harboring this modification. For this, we specifically developed a syngeneic mouse vaccination system where we had citrullinated a mouse cancer cell line, EMT6, and tested humoral responses in a normal, immunocompetent mouse. We found a robust antibody response that was unique to cells treated with active and not inactive PAD4. While this treatment induced a humoral response, it was not sufficient to block tumor growth of the EMT6 cells. This may be due to the fact that EMT6 is a very rapidly growing tumor and outpaced the ability of the humoral response to prevent rapid cancer cell proliferation, or that T-cell involvement is critical too.

In summary, these studies provide a simplified proteomics approach to identify protein substrates for PAD enzymes. These help to further our understanding of how PAD4 may promote tumor metastasis and induce large humoral responses to both extracellular matrix and cell surface proteins. We believe these new tools, methods and data sets should help the community to analyze, target and better understand citrullination by PAD4 in cells and *in vivo*.

## Methods

### Expression and purification of PAD4

C43 (DE3) Pro+, BL21 Gold (DE3), or BL21 ClearColi *E. coli* containing PAD4 expression vectors were grown in 2xYT at 37 °C to an OD-600 of 0.4–0.8 and then protein expression was induced by the addition of 0.5–1.0 mM IPTG. Incubation temperature was subsequently reduced to 18°C and the cultures were allowed to shake for 16–20 h. Cells were harvested by centrifugation and lysed using sonication. The lysate was centrifuged to remove inclusion bodies. The enzymes were purified by Ni-NTA resin with 0.5 mM TCEP supplemented to all buffers to prevent PAD4 oxidation. The purified enzyme was buffer exchanged to 50 mM Tris (pH 8), 400 mM sodium chloride, and supplemented with 0.5 mM TCEP. Purification steps were performed on ice to maintain high PAD4 enzymatic activity. Purified enzyme was aliquoted and flash frozen.

### Citrullinated histone H3 PAD4 activity assay

PAD4 activities in the absence or presence of antibodies were also assessed using a citrullinated histone H3 Western assay. 10-100 nM recombinant PAD4 were mixed with antibodies with various concentrations of calcium at 4°C for 45 min, and then incubated with 760 nM recombinant histone H3.1 (New England Biolabs). The reaction was incubated at 37°C for 110 min, followed by western analysis using an anti-citrullinated H3 primary antibody (Abcam Ab5103) and an anti-rabbit HRP secondary antibody. The reactions were then incubated at 37°C for 110 min and subsequently analyzed by western blot using an anti-citrullinated H3 antibody (Abcam, Ab5103) and anti-rabbit HRP secondary antibody. Images were acquired in Image Lab (v5.0) and processed with Image Studio Software (v5.2). IC_50_ measurements were obtained with technical triplicates and quantified using Fiji.

### Modified citrulline western blot assay

Lysate was harvested from Expi293T cells using a 1x RIPA + protease inhibitor solution. Antibody-PAD4 complexes were pre-formed at 4°C for 45 min. The complexes were then incubated at 37°C for 110 min and subsequently analyzed by western blot using an anti-modified citrulline detection kit (EMD Millipore). PVDF membrane was incubated with anti-citrulline probe overnight, then blocked and stained with a primary anti-modified citrulline antibody and secondary HRP linked anti-IgG antibody. HRP signal was detected using a BioRad ChemiDoc imager.

### Biolayer interferometry

BLI measurements were made using an Octet RED384 (ForteBio) instrument. Biotinylated PAD4 was immobilized on optically transparent streptavidin biosensors (ForteBio) and loaded until a 1 nm signal was achieved. After blocking with 10 μM biotin, purified binders in solution were used as the analyte. TBSTB was used for all buffers. Data were analyzed using the ForteBio Octet analysis software, and kinetic parameters were determined using a 1:1 monovalent binding model.^71^

### On-cell binding of PAD4 by flow cytometry

Adherent cells were lifted from culture with Versene (Fisher Scientific) at 37 °C, harvested by centrifugation (500 rcf), and resuspended in PBS in the presence of 1 uM PAD4. Cells were incubated at 4°C for 30 min, washed three times with cold 1% BSA in PBS (500 rcf, 5 min), and incubated on ice for 30 min with an anti-PAD4 primary antibody (Abcam) at a 1:500 dilution in 1% BSA in PBS. Cells were washed three times with cold 1% BSA in PBS (500 rcf, 5 min) then incubated with anti-rabbit IgG AlexaFluor 647 conjugated secondary antibody (Abcam) and incubated for 30 min at 4°C. Cells were washed three times with cold 1% BSA in PBS then incubated for 15 min with Propidium Iodide Ready Flow (Thermo Scientific) live/dead cell stain. Cells were analyzed on a Beckman Coulter CytoFLEX flow cytometer.

### On-cell activity of PAD4 and ZHER2-PAD4 fusion proteins

EMT-6 hHER2+ cells were harvested with Versene and harvested by centrifugation (500 rcf) and resuspended in DMEM with 2 mM Ca^2+^ and 0.5 mM TCEP supplemented. Cells were incubated for 60 min at 37 °C, rotating, in the presence of PAD4 fusion proteins, inactive enzyme controls, or vehicle. Cells were then washed three times with PBS and fractionated and lysed using a subcellular fractionation kit. Protease inhibitor (Sigma) and benzonase nuclease (Sigma Aldrich) were added to the lysate buffer. Membrane fractions were isolated and clarified by centrifugation (21000 rcf, 15 min) and protein concentration was quantitated by RapidGold BCA (Thermo Scientific). Normalized concentrations of lysates were separated by SDS-PAGE and transferred to PVDF membranes, then blotted for citrullination using the modified citrulline western blot kit (described above).

### Mouse immunization experiments

EMT6 hHER2+ cells were harvested with Versene and harvested by centrifugation (500 rcf) and resuspended in DMEM with 2 mM Ca^2+^ and 0.5 mM TCEP supplemented. Cells were incubated for 60 min at 37 °C, rotating, in the presence of PAD4 fusion proteins, inactive enzyme controls, or vehicle. Cells were then washed three times with PBS and then resuspended at 1e7 cell per mL in PBS and killed by heat-shock at 47 °C for 1 hour with shaking at 900 rpm. Heat-killed lysates were immediately flash-frozen until use.

BALB/c mice received subcutaneous injections of 100 µL heat-killed lysate in alternating flanks weekly for four weeks. At the end of the four weeks, mice were euthanized according to standard protocols per the Institutional Animal Care and Use Committee at the University of California, San Francisco. Blood and spleens were harvested for later use. Serum was separated from whole blood by centrifugation. Serum was then aliquoted and flash-frozen in liquid nitrogen. Frozen aliquots were stored at -80 °C. Splenocytes were harvested by homogenizing spleens using a 45 μm nylon mesh filter and the rubber end of a 3 mL syringe plunger in 2 mL PBS. The dissociated cells were harvested by centrifugation (600 rcf, 5 min). Pellets were resuspended in 2 mL ACK lysis buffer (Fisher Scientific) and incubated at room temperature for 3 min, at which point 10 mL PBS was added to neutralize the lysis. Cells were pelleted by centrifugation (600 rcf, 5 min), washed with 10 mL PBS again, and resuspended in 1 mL cryopreservation medium (Biolife) for gradual cooling and long-term storage in liquid nitrogen vapor phase.

### Serum flow cytometry

EMT-6 hHER2+ cells were harvested with Versene and harvested by centrifugation (500 rcf) and resuspended in DMEM with 2 mM Ca^2+^ and 0.5 mM TCEP supplemented. Cells were incubated for 60 min at 37 °C, rotating, in the presence of PAD4 fusion proteins, inactive enzyme controls, or vehicle. Cells were washed three times with cold 1% BSA in PBS (500 rcf, 5 min) and incubated on ice for 30 min with diluted serum (1:50) from immunized animals. Cells were washed three times with cold 1% BSA in PBS (500 rcf, 5 min) and incubated on ice for 30 min with goat anti-mouse IgG AlexaFluor 647 conjugate (Thermo) at a 1:200 dilution in 1% BSA in PBS. Cells were washed two times with cold 1% BSA in PBS (500 rcf, 5 min), once with PBS (500 rcf, 5 min), and incubated for 15 min in PBS with Propidium Iodide Ready Flow (Thermo Scientific) live/dead cell stain. Cells were analyzed on a Beckman Coulter CytoFLEX flow cytometer.

### Mouse immunization and challenge experiments

Mice were immunized as before. EMT-6 hHER2+ cells were harvested with Versene and harvested by centrifugation (500 rcf) and resuspended in DMEM with 2 mM Ca^2+^ and 0.5 mM TCEP supplemented. Cells were incubated for 60 min at 37 °C, rotating, in the presence of PAD4 fusion proteins, inactive enzyme controls, or vehicle. Live cells were then resuspended at 1e7 cell per mL in PBS and injected subcutaneously into the flanks of immunized or naïve BALB/c mice. Tumor growth and take rate was monitored periodically.

### Preparation of samples for mass spectrometry proteomics

Recombinant histone H3.1 protein and hPAD4 were incubated at 37 °C with TCEP and Ca^2+^ to generate citrullinated protein. Samples were then alkylated and digested on-bead using Preomics iST 96x kits (Preomics) and purified according to manufacturer’s protocols. Peptides were dried *in vacuo* and resuspended with 0.2% formic acid in before quantification with a Pierce Colorimetric Peptide Quantification Kit (Thermo). For cell lysate experiments, EMT6 and EMT6 hHER2+ cells were grown to confluency and lifted with Versene. Cells were washed once with ice cold PBS, then lysed with RIPA buffer supplemented with protease inhibitor. Lysates were spun down at max speed for 10 minutes at 4 °C and finally treated with PAD4 to generate citrullinated proteins in the lysate. Lysates were prepared for mass spectrometry analysis as described above.

### Off-line fractionation of mass spectrometry peptide samples

High-pH peptide fractionation was performed on an Agilent 1260 Infinity BioInert LC with an automated fraction collector. A 20-min method was performed on a Waters XBridge, Peptide BEH C18, 3.5 µm, 130 Å, 4.6 mm × 150 mm column with a flow rate of 800 µL/min. Mobile phase A (MPA) was 10 mM ammonium formate (Sigma-Aldrich, LC-MS grade), pH 10 and MPB was 20% 10 mM ammonium formate (pH 10)/80% methanol (Optima LC/MS grade, Fisher Scientific). The gradient ramped from 0 to 35% B from 0 to 2 min, 35 to 75%B from 2-8 min, 75% to 100% B from 8 to 13 min, followed by washing at 100% B from 13 to 15 min and equilibration at 0% B from 15 to 20 min. UV absorbance at 210 and 280 nm was recorded. To generate deep citrullination libraries from peptides of EMT6 cells, 16 fractions were collected from 5 to 18 min and concatenated into the final fractions by combining fraction 1 and 9, fraction 2 and 10, etc., resulting in 8 final fractions. Samples were dried down in a GeneVac prior to being resuspended in 0.2% formic acid (Fisher Scientific, LC-MS grade) for LC-MS analysis.

### Mass spectrometry proteomics of citrullinated samples

Samples were analyzed using a Vanquish Neo UHPLC system coupled to an Orbitrap Eclipse mass spectrometer (Thermo Fisher Scientific, San Jose, CA). Resuspended peptide samples were loaded onto a 60 cm long 75 *µ*m inner diameter column (Ion Opticks, Aurora Ultimate). Mobile phase A was composed of water and 0.2% formic acid (FA) while mobile phase B was composed of 80% ACN and 0.2% FA. Separation was performed using a gradient elution of 7-48% mobile phase B over 65 minutes followed by 48-60% mobile phase B over 7 minutes and 99% mobile phase B for 24 minutes. Flow rate was held at *300 n*L/min. MS^1^ survey scans of peptide precursors from 350 – 2000 *m/*z were performed at a resolving power of 60,000 with an AGC target of 1 x 10^6^, and maximum injection time of 50 ms. For the HCD-pd-EThcD method, a neutral loss of 43.0058 *m/z* ± 10 ppm from the precursor was listed to trigger a subsequent EThcD fragmentation. The HCD MS/MS scans were set to an AGC target of 1.25 x 10^5^, a resolving power of 30,000, and maximum injection time of 100 ms. The ETD reaction time was set to an AGC Target of 1.25 x 10^5^, a resolving power of 60,000, and a maximum injection time of 118 ms.

### Mass spectrometry data analysis

Thermo .RAW files were analyzed in FragPipe (v21.1) against a FASTA file containing the complete murine proteome with isoforms plus human PAD4, human HER2, and common contaminant proteins (downloaded from UniProt April 30, 2024). with trypsin selected as the enzyme and 4 missed cleavages allowed.^72–75^ Cysteine carbamidomethylation (+57.02146) was fixed, while methionine oxidation (+15.9949 Da), protein N-terminal acetylation (+42.0106 Da), and arginine citrullination (+0.984 Da) were chosen as variable modifications. A localization threshold of 0.75 was used for citrullination. Peptide spectral matches (PSMs) identified which ended in a C-terminal citrulline residue were filtered out of the dataset as the neutral citrulline residue should induce a missed tryptic cleavage. MS^1^ and MS/MS mass tolerances were set to ±20 ppm. All other parameters were set as default.

## Data Availability

The LC-MS/MS data including .RAW, .mzML files and modified peptide identifications have been uploaded to the MassIVE proteomics data repository (PXD072496).

## Acknowledgments

We thank Drs. Kevin Leung and Jamie Byrnes for helpful assistance with proteomics discussions and the Wells Lab broadly for helpful discussions and expertise. J.A.W. was supported by generous grants from NIH NCI1P41CA196276, CA191018, and NIH GM097316 and commercial funding from Bristol Myers Squibb. Animal studies were supported by the HDFCCC Shared Resource Facility, Preclinical Therapeutics Core, through NIH (P30CA082103). T.M.P.C. was supported by the National Cancer Institute of the National Institutes of Health through a F32 Postdoctoral Fellowship (F32CA298768). C.S.D is an AP Giannini Foundation Postdoctoral Fellow.

