## Supplementary material for "Cellular consequences, citrullination substrates, and antigenicity resulting from wild-type and targeted PAD4 on cell surfaces": Fig S

**Authors & Affiliations**

Sophie Kong^1,2 ‡^, Trenton M. Peters-Clarke^1‡^, Corleone S. Delaveris^1,3^, Paul Phojanakong^4^, Veronica Steri^4^, James A. Wells^1,4*^

*^1^Department of Pharmaceutical Chemistry, University of California San Francisco, San Francisco, California, 94158, USA.*

*^2^Current address: Epi Biologics, San Mateo, CA 94403 USA.*

*^3^Current address: Inversion Therapeutics, San Francisco, California, 94158, USA.*

*^4^Preclinical Therapeutics Core, University of California San Francisco, San Francisco, California, 94158, USA.*

*^5^Department of Cellular & Molecular Pharmacology, University of California San Francisco, San Francisco, California, 94158, USA.*

^‡^*Authors contributed equally*

**Figure S1.** Schematic of transwell migration assay	S2

**Table S1.** Top citrullinated protein hits identified from proteomic analysis	S3

**Figure S2.** Global heat map of citrullinated proteins mapped to known HER2 interactors	S4

**Figure S3.** Tumor volume measured over time of subcutaneously implanted EMT6 tumors following vaccination.	S5

**Figure S4.** Upset plot displaying the number of common and unique peptides across conditions	S6

**Figure S5.** Tumor regression studies of mice implanted with EMT6 hHER2+ cells	S7

**Figure S6.** Breakdown of scan types within HCD-pd-EThcD for citrullination profiling of EMT6 lysate	S8

**Figure S7.** Gene ontology enrichment for citrullinated proteins targeted by ZHER2-PAD4 within EMT6 lysate	S9

**Figure S8.** Gene ontology enrichment for citrullinated proteins targeted by ZHER2-PAD4 within the surfaceome of SKBR3 cells	S10

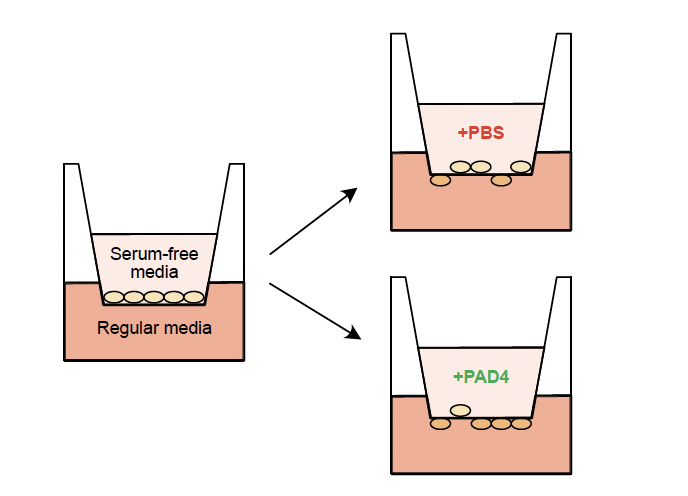
 **Supp Fig. S1. Schematic of transwell migration assay measuring the count of cells migrating from serum-free media (upper chamber) into the regular media (lower chamber).**

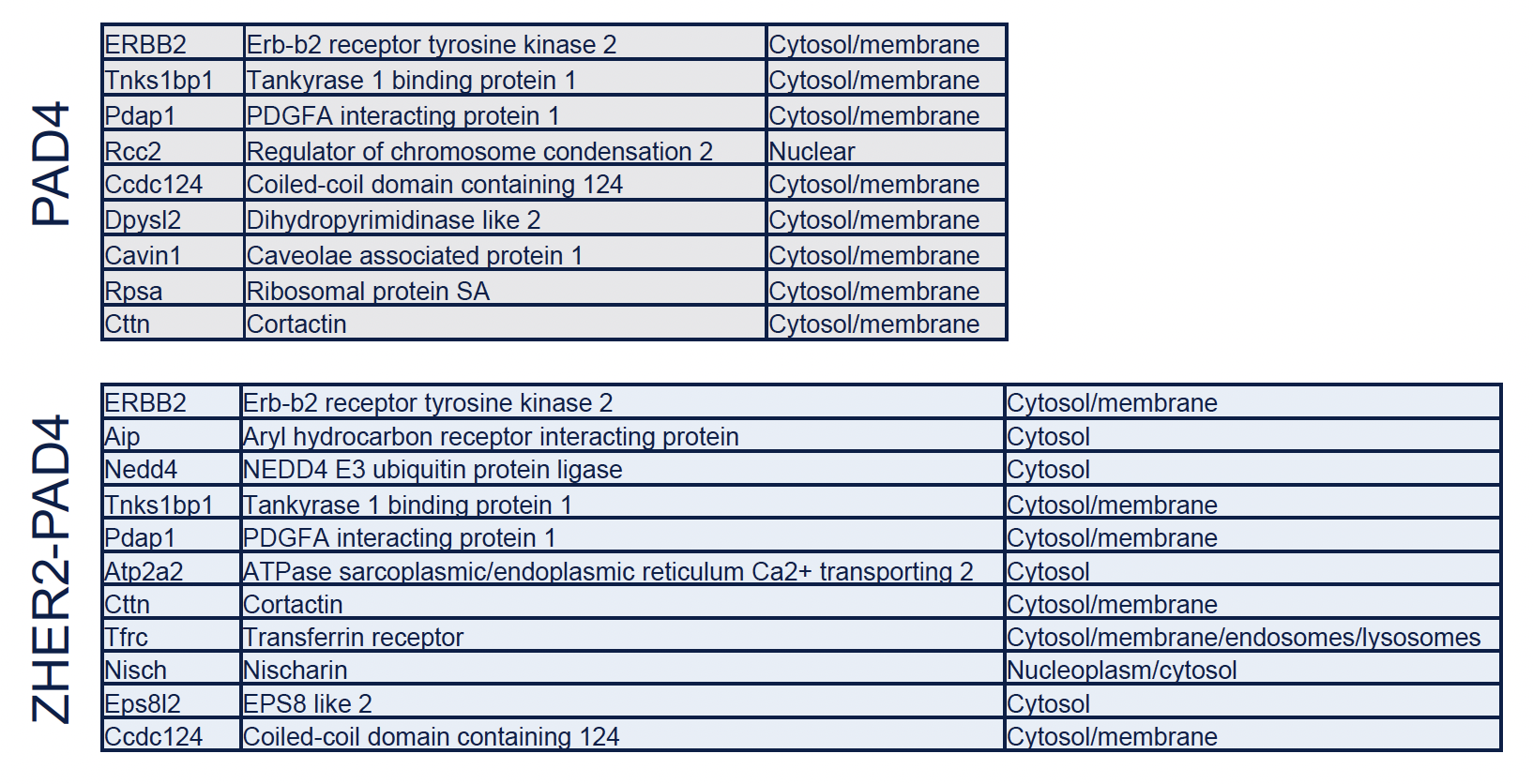
 **Supplemental Table S1. Top citrullinated protein hits identified from proteomic analysis and their canonical location.**

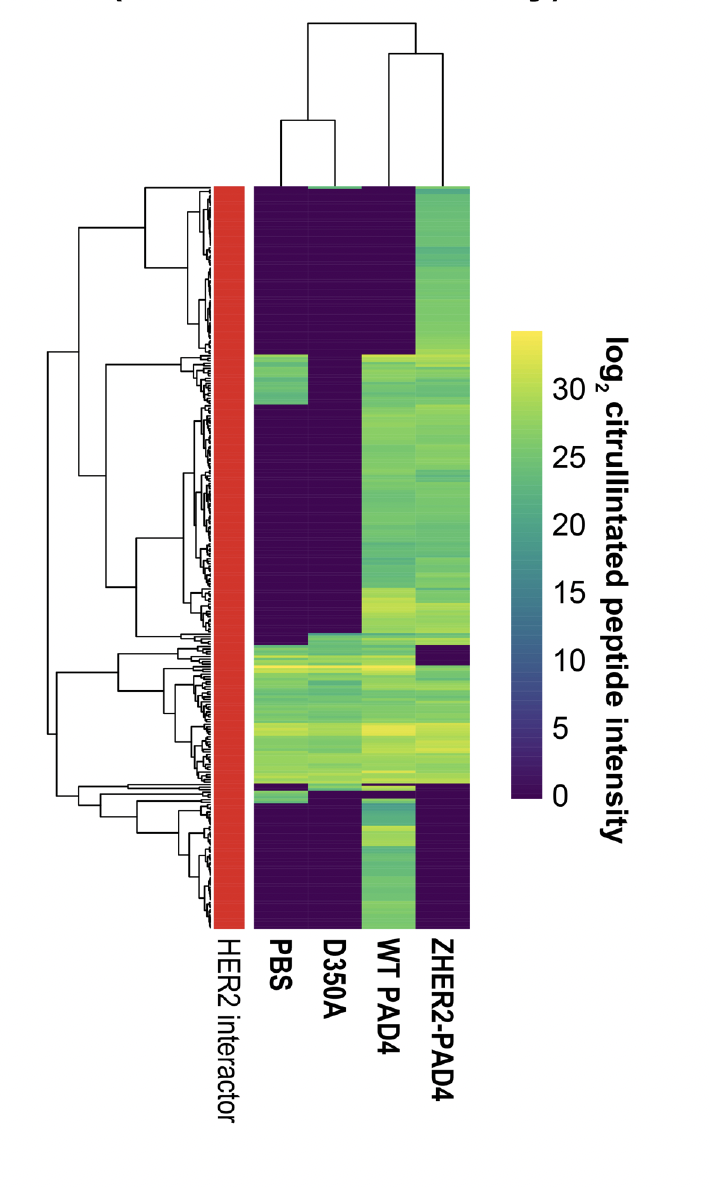

**Fig S2. Global heat map of citrullinated proteins mapped to known HER2 interactors .** A companion to Figure 4F, this heatmap highlights citrullination only for known HER2 interactors’ sites. While some background endogenous citrullination is observed across all four conditions, novel sites emerge for active PAD4 conditions.

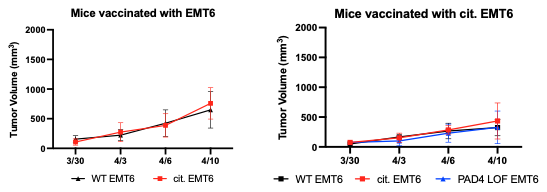

**Fig. S3. Tumor volume measured over time of subcutaneously implanted EMT6 tumors following whole cell vaccination.** Black plot corresponds WT EMT6 tumors implanted following vaccination, red corresponds to citrullinated tumors, and blue corresponds to tumor cells treated with inactive PAD4.

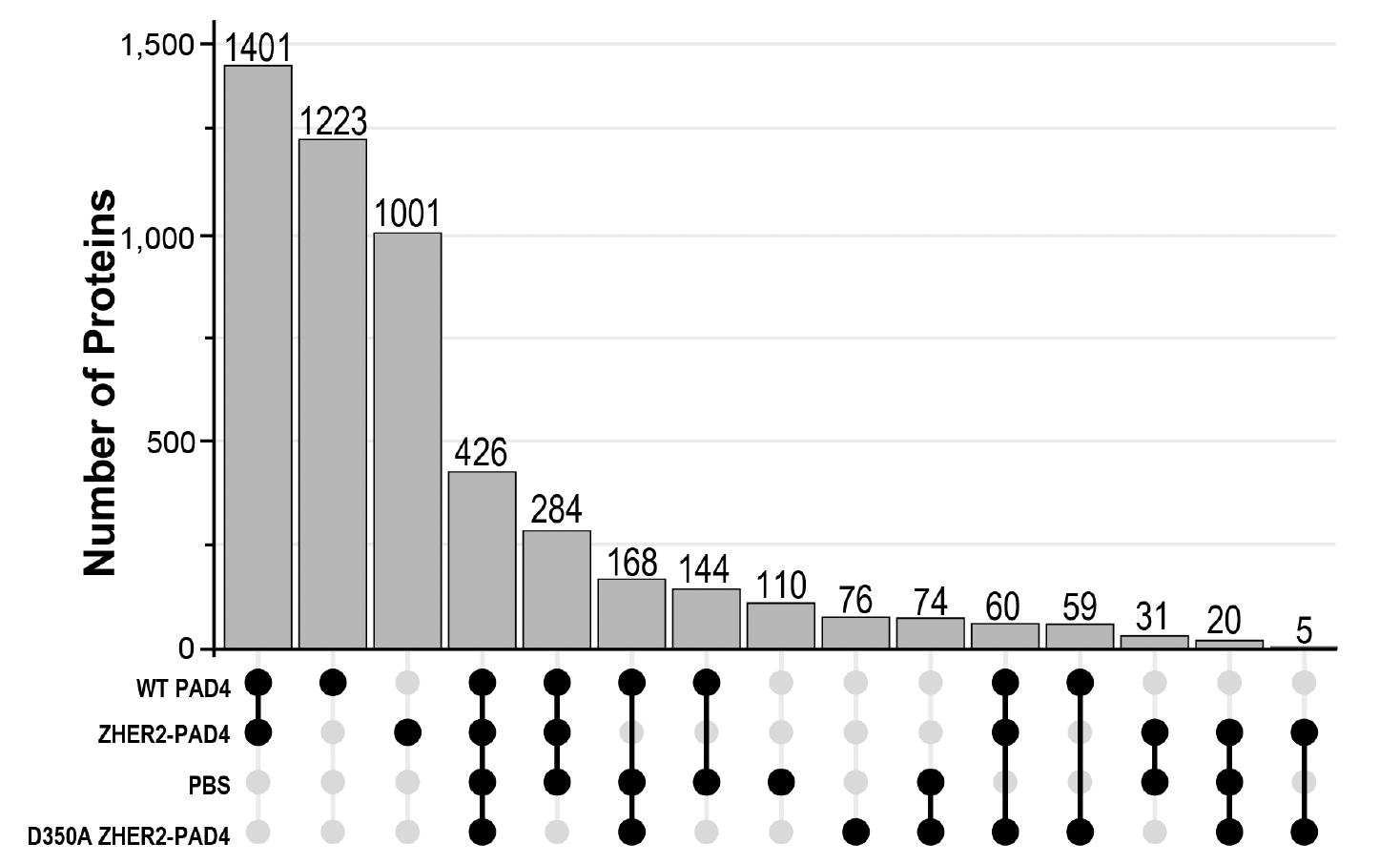
 **Fig S4. Upset plot displaying the number of common and unique peptides after EMT6 cell lysate was treated under indicated conditions.** The most similarity is seen for active PAD4 conditions, while WT PAD4 and ZHER2-PAD4 treated lysates produce many unique citrullinated protein substrates.

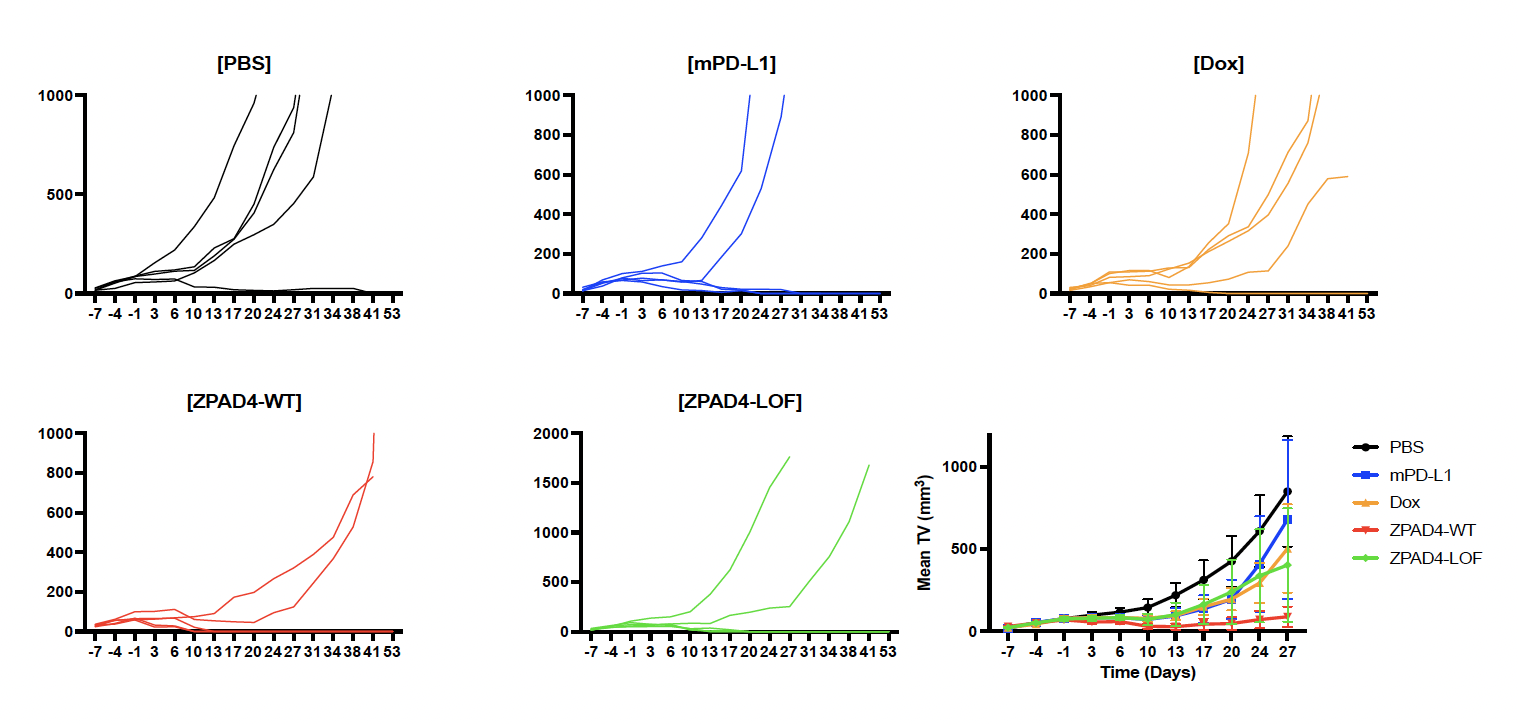
 **Fig. S5. Tumor regression studies of mice implanted with EMT6 hHER2+ cells.** Mice subcutaneously implanted with EMT6 hER2+ cells were treated with PBS, mPD-L1, Dox, ZPAD4-WT, or ZPAD4-LOF. Mice treated with ZPAD4-WT had an observed delay in tumor growth compared to negative control and even positive control groups. ß

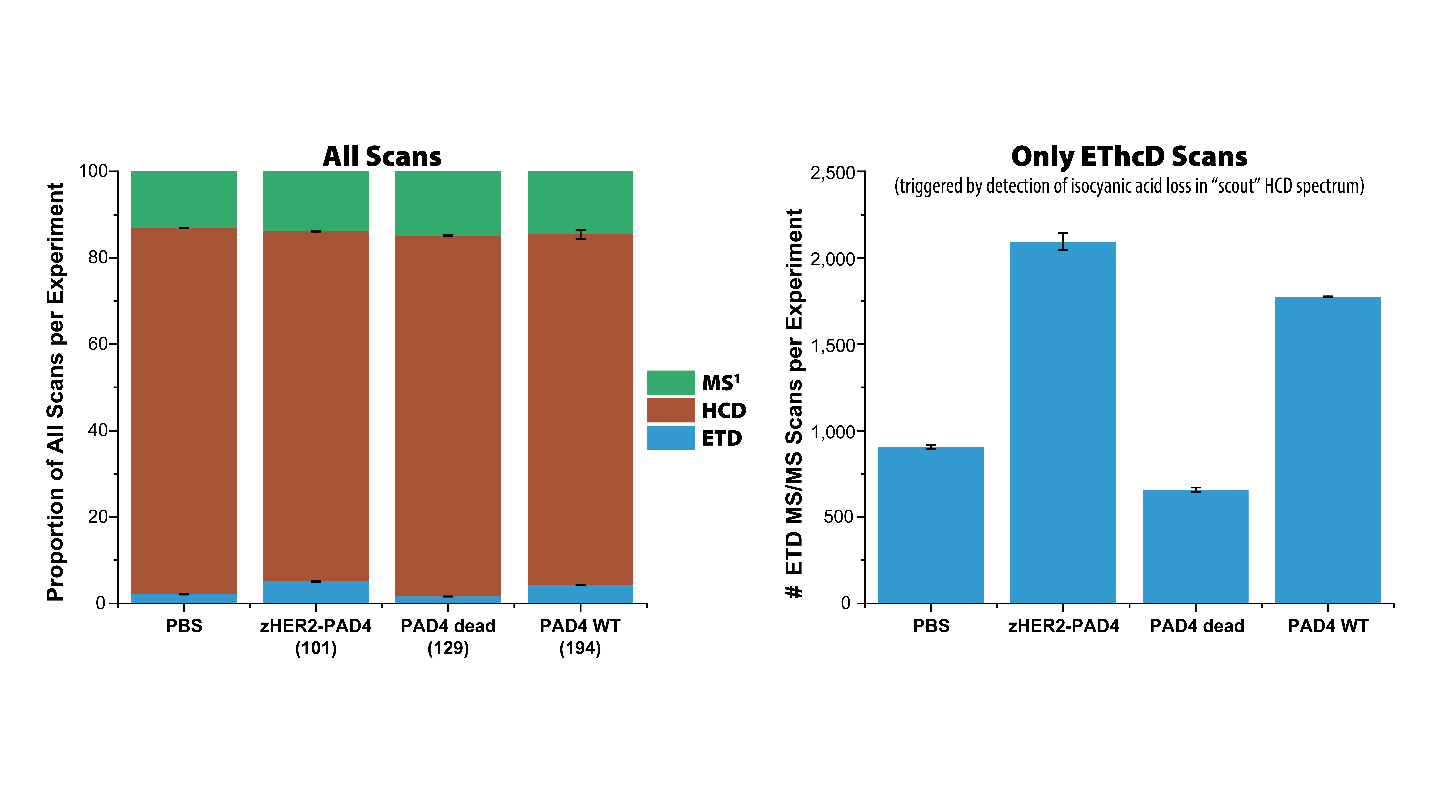
 **Fig S6. Breakdown of scan types within HCD-pd-EThcD intelligent data acquisition for citrullination profiling of EMT6 lysate.** Triggering an EThcD scan necessitates longer scan times and slower duty cycles. Performing a quick HCD scan will release a labile isocyanate neutral loss from citrulline residues, an event that produces a unique diagnostic ion. Matching this mass shift (-43 Da) from the intact precursor (±10 ppm) in the scout HCD scan triggers a sequence informative and citrulline localizing EThcD scan. While all conditions produced some level of EThcD scans, the relative number of EThcD scans was highest for active PAD4 (WT and zHER2 targeted). The zHER2-PAD4 treatment of EMT6 lysate generated the highest number of triggered scans, over double the number of EThcD scans as the PBS and PAD4 LOF treatments. This indicates high enrichment of citrulline-containing peptides within the zHER2-PAD4 sample.

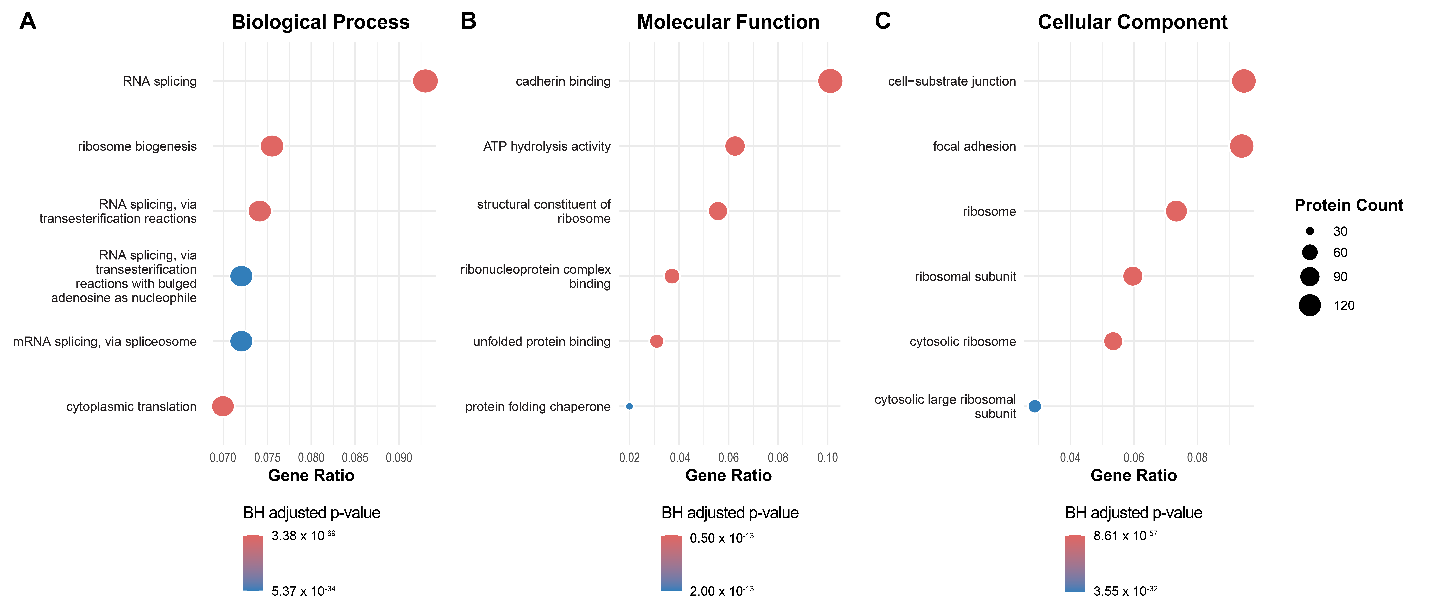
 **Fig S7. Gene ontology enrichment analysis for citrullinated proteins targeted by ZHER2-PAD4 within EMT6 lysate.** Proteins were enriched according to their biological processes, molecular functions, and cellular components. Circle size represents the number of proteins in that gene ontology term, while the color of the circle represents the Benjamini-Hochberg (BH) corrected p-value. Gene ratio refers to the fraction of all citrullinated proteins by ZHER2-PAD4 that were categorized into the associated GO term.

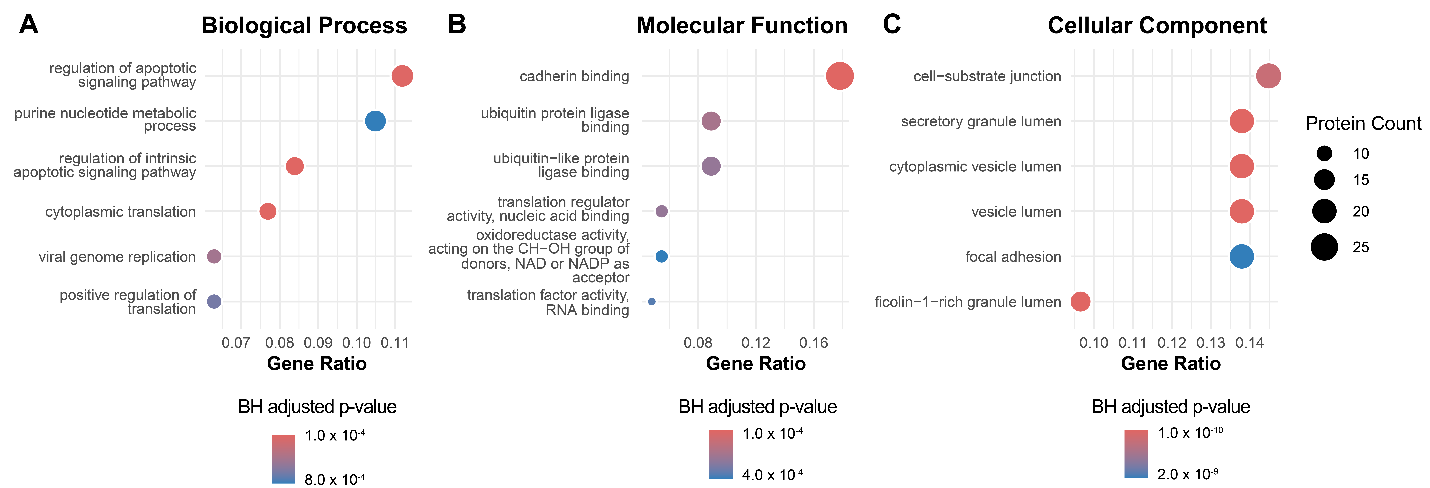
 **Fig S8. Gene ontology enrichment analysis for citrullinated proteins targeted by ZHER2-PAD4 within the surfaceome of SKBR3 cells.** Proteins were enriched according to their biological processes, molecular functions, and cellular components. Circle size represents the number of proteins in that gene ontology term, while the color of the circle represents the Benjamini-Hochberg (BH) corrected p-value. Gene ratio refers to the fraction of all citrullinated proteins by ZHER2-PAD4 that were categorized into the associated GO term.
